# Mapping Light-Induced Conformational Dynamics of Pigeon Cryptochrome 4 by HDX-MS: Structural Transitions from Spin Pair Formation to Activated Conformational States

**DOI:** 10.64898/2026.08.27.747556

**Authors:** Gargi S. Jagdale, Victoria Fan, Pankaj Dubey, Alan Pham, Emily Jiang, Anthony T. Iavarone, Judith P. Klinman

## Abstract

The navigational prowess of migratory birds is thought to arise from light-dependent radical-pair chemistry in cryptochrome 4 (CRY4), yet the slow structural transitions that couple photochemistry to signaling remain elusive. Here, we combine temperature-controlled steady-state UV–visible spectroscopy and hydrogen–deuterium exchange mass spectrometry (HDX-MS) to elucidate the photochemical and conformational dynamics of pigeon CRY4 (*Cl*CRY4). Steady-state measurements at 5–25 °C reveal that lower temperatures slow FAD photoreduction and prolong the FADH^•^ signaling state. This occurs without a solvent kinetic isotope effect, implicating a conformational change rather than proton transfer as the rate determining step in FADH^•^ formation. Simultaneous HDX-MS under blue-light exposure identifies protection near the FAD-binding site and C-terminal region. To enhance sensitivity, we developed a pump-probe HDX-MS approach at 10 °C. This reveals eight peptides (within the phosphate-binding loop, protrusion motif, electron-transfer-chain loops and C-terminal tail) that exhibit rapid (≤10 s) and sustained light-induced protection, delineating early conformational rearrangements as a prerequisite for FADH^•^ accumulation. The findings of slower onset HDX protection as well as a bimodal pattern of deuterium uptake in the phosphate-binding loop further identify a local redistribution of conformational substates on the time scale of the accumulation of the signaling species FADH^•^ . Site specific mutagenesis within the CTT supports the findings, which lead to a model in which blue light triggers rapid clamping down of protein near the two regions of spin pair separation, followed by a rate limiting closure of a surface loop. The resolution of time-dependent structural transitions that follow photoactivation of CRY4 resolves the interface between quantum radical-pair formation and classical conformational changes, while providing an enhanced structural framework for the molecular events that underlie avian magnetoreception.

## Introduction

The ability of migratory birds to navigate across vast distances with remarkable precision has long intrigued scientists and the public alike. A growing body of evidence supports the existence of a specialized sensory system that enables birds to use the Earth’s magnetic field in orientation and migratory behavior -a phenomenon known as magnetoreception (1,2). Two principal models have been proposed to explain avian magnetoreception: the radical pair mechanism (RPM) and the magnetite-based model. The RPM posits that photoinduced electron transfer reactions in cryptochrome (CRY) proteins generate spin-correlated radical pairs whose recombination kinetics are sensitive to weak magnetic fields, thereby providing directional information (3–5). In contrast, the magnetite-based model suggests that magnetite crystals within specialized sensory cells act as biological compass needles, transducing magnetic field information via mechanical or ion channel-mediated pathways (6,7).

Among these, the RPM has gained substantial support, particularly due to its ability to explain the light dependence and inclination sensitivity of the avian magnetic compass (8,9). Cryptochromes are evolutionarily conserved flavoproteins that function as blue/UV-light photoreceptors in both plants and animals and are closely related to DNA photolyases (10,11). In birds, cryptochrome 4 (CRY4) has emerged as the leading molecular candidate for the primary magnetoreceptor. CRY4 is highly expressed in the retina, especially in the outer segments of double cones and long-wavelength single cones cell types that are implicated in magnetoreception (12,13). Notably, CRY4 expression exhibits seasonal regulation and peaking during migratory periods, which further supports its role in navigation (14).

Recent biochemical and spectroscopic studies have demonstrated that avian CRY4 proteins, including those from pigeon (*Cl*CRY4), chicken (*Gg*CRY4), and European robin (*Er*CRY4), undergo light-driven electron transfer reactions that generate long-lived FADH^•-^ and Trp^•+^ radical pairs (15–18) that are sensitive to geomagnetic fields.

Detailed mechanistic studies, including those by Xu et al. (19), have elucidated the time scales and sequence of photochemical steps leading to the signaling state in avian CRY4. Upon absorption of UV or blue light, the FAD cofactor is excited (FAD*), triggering a rapid (picosecond) electron transfer from the closest tryptophan residue to FAD, resulting in the formation of a singlet radical pair (^1^[FAD^•–^ + TrpH^•+^]), **Figure 1c**. The distance between the radical pair increases as the electron transfer takes place from subsequent Trp residues (picosecond to nanosecond). This radical pair can interconvert with its triplet state (^3^[FAD^•–^ + TrpH^•+^]) on the nanosecond to microsecond timescale due to hyperfine and Zeeman interactions that are modulated by the geomagnetic field. The fate of the radical pair is determined by two competing processes: (1) spin-selective back electron transfer to the ground state (FAD_ox_), or (2) proton transfer (>100 μs), which neutralizes and stabilizes the radical anion to form FADH^•^ as the proposed signaling state. Notably, Xu et al. showed that the electron transfer steps through the tryptophan chain occur at rates >10^10^ s^−1^, while the formation and decay of the stabilized FADH^•^ state occur on much longer timescales. The dynamic equilibrium between different radical pair states and the branching between recombination and stabilization are predicted to be finely tuned to maximize both magnetic sensitivity and signaling efficiency in migratory birds.

**Figure 1.**
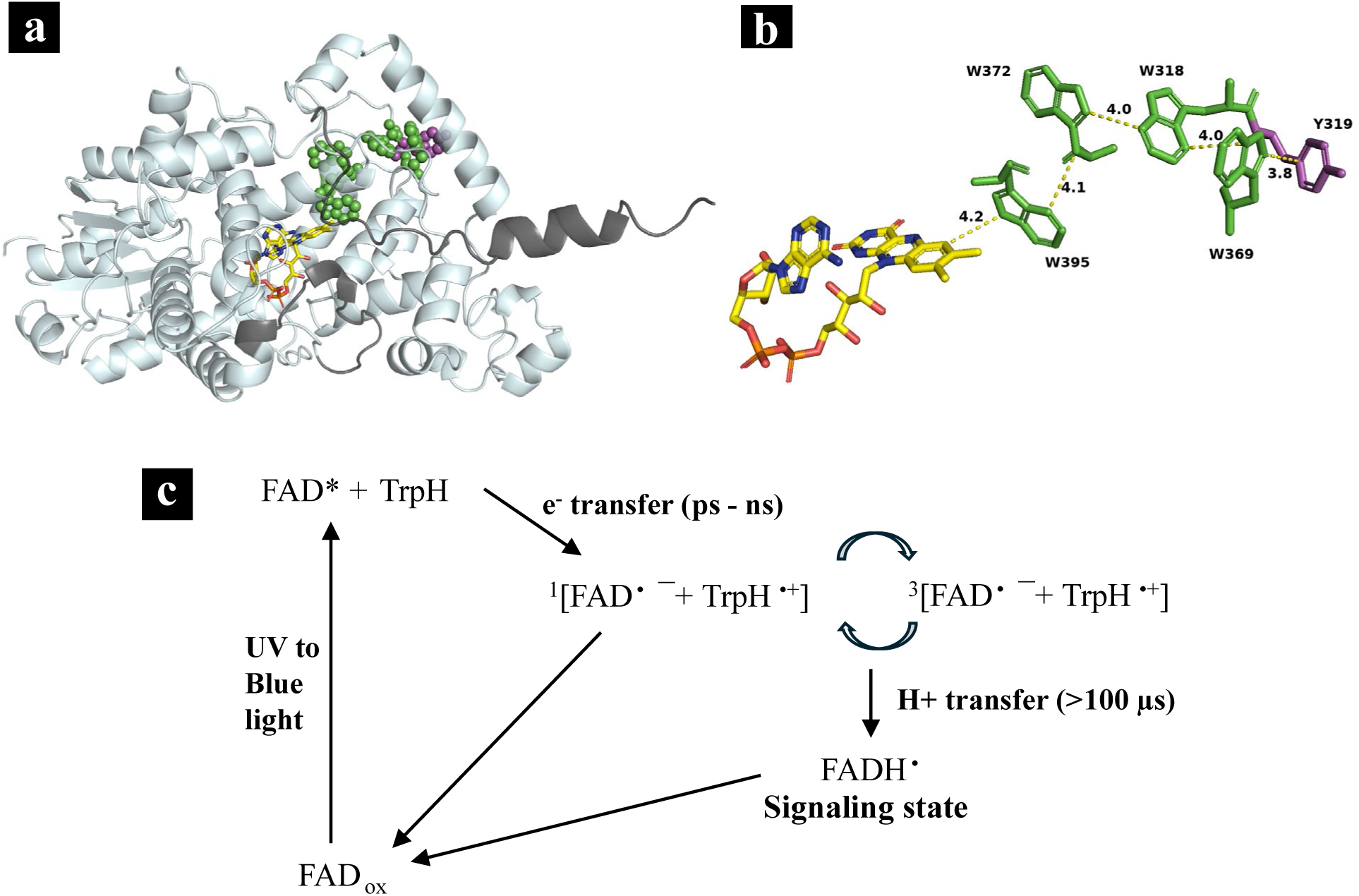
Structure and photochemistry of pigeon cryptochrome 4 (*Cl*CRY4). **(a)** AlphaFold model of *Cl*CRY4 (AF-A0A386QUR4-F1-v4). Regions unresolved in the available crystal structure (PDB 6PU0) are shown in dark gray: the phosphate-binding region (residues 229–243) and the C-terminal tail (CTT, residues 498–525). The FAD cofactor (yellow sticks) was modeled from the crystal structure; the tryptophan tetrad is shown as green spheres and the surface-exposed tyrosine Y319 as magenta spheres. **(b)** Stick representation of the electron-transfer chain showing the FAD cofactor (yellow), the tryptophan tetrad (W395, W372, W369, W318; green), and the candidate fifth redox-active residue Y319 (magenta); edge-to-edge distances (Å) between adjacent partners are indicated. **(c)** Simplified scheme of avian CRY4 photochemistry. Absorption of UV/blue light promotes FADox to its excited state (FAD*), triggering picosecond-to-nanosecond electron transfer from the proximal tryptophan to form the singlet radical pair 1[FAD•– + TrpH•+], which interconverts with its triplet state 3[FAD•– + TrpH•+]. Sequential electron-hole migration along the tryptophan tetrad (W395 → W372 → W369 → W318) progressively increases the separation between radical-pair partners, slowing back electron transfer and extending the lifetime of the radical pair. Proton transfer (>100 µs) stabilizes the radical to form the neutral semiquinone FADH•, the proposed signaling state; spin-selective back electron transfer regenerates FADox. Scheme adapted from refs (15,19).

Structural and computational analyses have revealed that avian CRY4s possess an extended tryptophan tetrad, a potential fifth redox species Tyr residue, and unique active site residues that facilitate efficient electron transfer and radical pair stabilization (15,17,20). For example, *Cl*CRY4 and *Gg*CRY4 both display a red-shifted FADH^•^ absorption maximum and unusually prolonged radical lifetimes compared to plant and insect cryptochromes, suggesting evolutionary adaptation for magneto-sensory function (15,17,21). The crystal and AlphaFold-based structures of *Cl*CRY4 used throughout this work locate the FAD cofactor, the tryptophan tetrad (W395, W372, W369, W318), and the surface-exposed Y319 within the photolyase-homology region and resolve the phosphate-binding loop and C-terminal tail as the principal mobile elements (**Figure 1a**). The corresponding electron-transfer chain, in which sequential hops connect the FAD to the four tryptophans and the candidate fifth redox residue Y319, is shown in **Figure 1b**.

Despite these advances, the molecular mechanisms by which CRY4 activation is coupled to downstream signaling pathways have remained poorly understood. In the radical pair model, it is hypothesized that light-induced electron transfer and subsequent conformational changes in CRY4 are transduced into a cellular signal, and the nature of this signal and its transmission to the nervous system are active areas of investigation (4,9). Recent studies suggest that the C-terminal tail (CTT) of CRY4 may play a critical role in this process. Light-induced conformational changes in the CTT and adjacent regions have been observed in both pigeon and chicken CRY4, as well as in other cryptochromes, and are thought to modulate interactions with signaling partners (15,17,21). In chicken CRY4, for instance, antibody accessibility assays and immunoprecipitation experiments have shown that CTT undergoes light-dependent shielding, suggesting a dynamic structural role in signal propagation (21). In a related fish CRY4, conserved electron transfer pathways have been implicated in similar signaling processes (22). These dynamic structural rearrangements are thought to be essential for coupling the initial photochemical event to the activation of intracellular signaling cascades. Similarly, molecular dynamics simulations of *Cl*CRY4 indicate that light-induced rearrangements in the phosphate-binding loop and CTT could facilitate the recruitment of downstream effectors, with the C-terminal region showing rapid response to photoactivation and sustained increases in mobility coupled with tryptophan rearrangements (17).

Several models have been proposed for downstream signaling in avian magnetoreception. One prominent hypothesis suggests that photoactivated CRY4 interacts with cone-specific G proteins, initiating a signaling cascade analogous to classical phototransduction in visual pigments (23,24). Surface plasmon resonance and co-immunoprecipitation studies have provided evidence for direct, light-dependent binding between CRY4 and Gα subunits in the retina (25). Alternatively, CRY4 has been proposed to form complexes with magnetite-associated proteins such as MagR, potentially acting as a biocompass through protein-protein interactions and mechanical coupling (23). However, the physiological relevance of these complexes remains to be fully established.

In this study, we present a detailed biophysical and structural analysis of pigeon CRY4 (*Cl*CRY4), focusing on directly correlated time- and temperature-dependent photochemistry and light-induced conformational dynamics. Using steady-state UV-visible spectroscopy and hydrogen-deuterium exchange mass spectrometry (HDX-MS), we demonstrate that *Cl*CRY4 exhibits robust temperature-coupled photochemistry and rapid, localized structural rearrangements upon blue-light activation. We identify the CTT, the loop containing the potentially redox active Tyr residue adjacent to the Trp quartet, and the phosphate-binding loop as key dynamic regions. We note that analyses of HDX-MS at reduced temperature have allowed close to 100% isolation of the putative signaling species, capturing conformational changes that occur on a timescale either preceding or accompanying the high accumulation of FADH^•^. From our findings and those from studies of European robin, chicken and fish CRY4, as well as other cryptochromes and photolyases, a model for site-specific, light-induced conformational changes in CRY4 can be envisioned that ratchets down rapid, light-driven spin pair production toward a downstream hierarchy of increasingly slower conformational states Beyond its natural role, the same spin-correlated radical-pair chemistry that underlies CRY4’s magnetosensitivity is increasingly being explored as a platform for engineered, genetically encoded magnetic-field sensors,(26,27) and the kinetic and structural determinants defined here, including the temperature-tunable lifetime of the FADH• signaling state and the CTT clamp mechanism, may inform efforts to rationally tune such sensors(28).

## Results

### Steady State Photochemistry of *Cl*CRY4: Temperature Dependence

The temperature dependence of *Cl*CRY4 photochemistry was investigated using steady-state UV-vis absorption spectroscopy at 25, 18, 10, and 5 °C. The experiments were performed using *Cl*CRY4 protein at a concentration of 25–35 µM, pH 8 (See SI). Spectra were first recorded in the dark (dim red light), after which the protein was irradiated with 250 lux blue light (450 nm), and absorption spectra were collected every 30 seconds over a period of 20 minutes. The differential spectra after light excitation (vs dark) recorded over time is shown in **Figure S1**. The raw traces were analyzed to determine the concentrations of FAD redox species, FAD_ox_, FADH^•^ and FADH**^−^**over time, normalized to the total FAD concentration determined from the dark-state absorbance at 450 nm, where only FAD_ox_ is present, following the approach established by Zoltowski et al. (15) and described in detail in the SI.

At 25 °C, the photochemical behavior of *Cl*CRY4 is consistent with previous reports (15). A decrease in temperature from 25 °C to 5 °C leads to a slower reduction of FAD_ox_ (**Figure 2**, black trace) and a decrease in both the formation and decay rates of FADH^•^ (red trace). At 10 °C and 5 °C, after its maximum formation, FADH^•^ exhibited minimal decay over the experimental timescale (20 min), and only a small fraction of FADH**^−^** was detected. The fully reduced species is assigned as the anionic hydroquinone (FADH^−^), rather than the neutral hydroquinone (FADH_2_), on the basis of two considerations: the characteristic absorption of the fully reduced flavin near 380 nm matches that of the anionic form, and the pK_a_ of the flavin N1 position (≈ 6.7 for free flavin) dictates that the anion likely predominates at the experimental pH of 8 (>90% FADH^−^). This assignment is consistent with prior CRY4 and photolyase studies (15,18,29–31).

**Figure 2.**
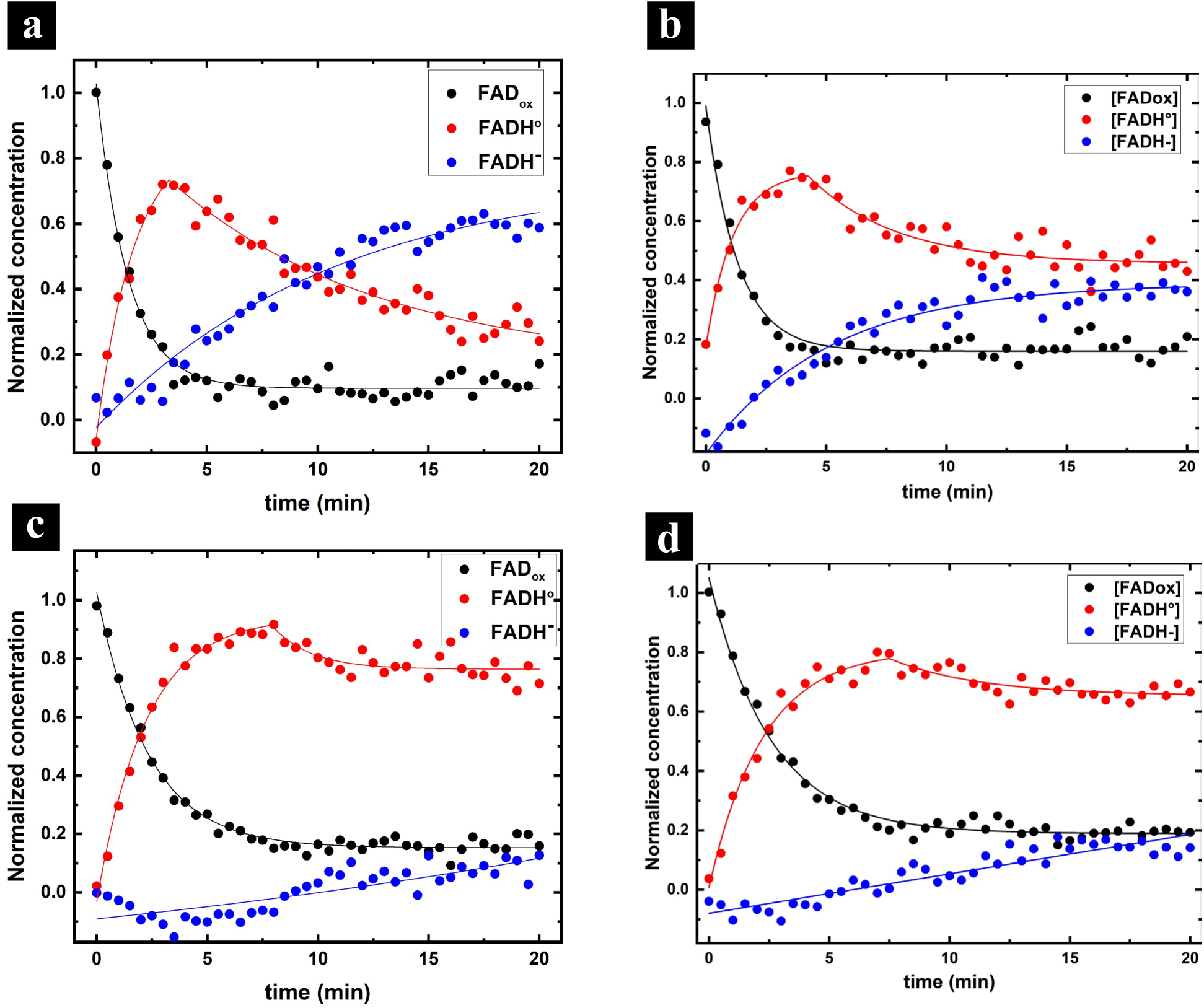
Temperature dependence of *Cl*CRY4 photochemistry monitored by steady-state UV-vis spectroscopy. UV-vis spectra were recorded every 30 s for 20 min under continuous 450 nm illumination (250 lux). Normalized concentrations of FADox (black), FADH• (red), and FADH– (blue), with respect to total FAD concentration, derived from the spectra as described in the SI, are plotted versus time at **(a)** 25 °C, **(b)** 18 °C, **(c)** 10 °C, and **(d)** 5 °C. Lower temperatures slow the decay of FADox and both the formation and decay of the proposed signaling state FADH•. At 10 °C and 5 °C the FADH• decay is very slow, allowing FADH• to accumulate with little formation of the third species (assigned to FADH–). Conditions near 10 °C therefore isolate and capture the signaling state (FADH•).

Previous studies have suggested that pH (or temperature) can influence CRY4 photochemistry, potentially due to changes in the protonation state (or pK_a_) of histidine residues that have pK_a_ values within the experimental pH range.(19) However, due to the lack of definitive information regarding the proton donor for FAD^•-^ protonation to FADH^•^ and the involvement of His residues specifically in *Cl*CRY4 photochemistry, the precise mechanistic basis for the temperature-dependent differences in photochemical reduction of FAD in *Cl*CRY4 remains unclear. To further probe the protonation step, steady-state photochemical analysis was conducted in D_2_O buffer at 25°C, pH 7.6 (pD = 8) (**Figure S2**). No significant differences were observed in the decay or formation kinetics of FAD_ox_ and FADH^•^ species in D_2_O vs H_2_O, ruling out proton transfer as the rate-limiting step in FADH^•^ formation. Importantly, these temperature-dependent photochemistry experiments define conditions under which the FADH• signaling state can be accumulated and stabilized, providing the experimental basis for the HDX-MS analysis below of the light-induced conformational changes proposed to underlie downstream cell signaling.

### Light-induced Structural Changes in *Cl*CRY4 Investigated by Hydrogen-Deuterium Exchange Mass Spectrometry

#### HDX-MS Analysis of ClCRY4 in the Ground State (Dark Conditions)

To characterize the structural properties of *Cl*CRY4 in its ground state, hydrogen-deuterium exchange mass spectrometry (HDX-MS) experiments were performed under dark conditions (dim red light) at 25 °C and 10 °C. The experimental protocol, described in detail in the Methods section of the SI, involves incubating the protein samples (5 µM final concentration) in 90% deuterated buffer, pH 7.6, across 14 time points, from 10 seconds to 20 minutes, followed by quenching, digestion and LC-MS analysis. Deuterium uptake was analyzed for 37 peptides (**Figure S3, Table S1**), achieving 98.86 % sequence coverage and an average peptide length of 12 residues. Two peptides, 323–335 and 336–342, were that were detected only at 25 °C and exhibited less than 5% deuterium uptake under dark conditions, with no significant change upon blue-light exposure (included in **Figure S5**). These results indicate specific regions that remain structurally protected and distinct from those involved in light-induced conformational changes.

The deuterium uptake profiles at selected time points (10 s, 1.5 min, 5 min and 20 min) at 25 °C and 10 °C are presented in **Figure S4**. Higher temperature correlated with greater deuterium uptake, reflecting increased protein dynamics. Additionally, within a given temperature, longer exposure to deuterium resulted in progressively higher uptake for each peptide. Complete time-dependent deuterium uptake traces under dark conditions for all peptides are shown (red) in **Figures S5** (25 °C) and **S6** (10 °C). Under these conditions, *Cl*CRY4 exhibited low to moderate D-uptake, indicating largely a solvent-protected and well-structured protein. However, two regions, one proximal to the FAD cofactor and another at the C-terminal, displayed >50 % deuterium uptake, indicating greater flexibility and solvent exposure.

The %D uptake after 5 minutes of exchange at 10 °C - the probe time used in subsequent experiments (See *Pump-Probe HDX-MS*) - is shown in **Figure 3a**. Most regions of the protein exhibited less than 50 % uptake, except for the FAD-interacting site (part of the phosphate binding loop, peptide 229-243) and the C-terminal region (peptide 404-412, and CTT peptides 496-525). Notably, the CTT peptides 496-506 and 506-525 displayed rapid saturation in the D-uptake, indicative of a highly flexible and solvent-exposed region. Interestingly, there is no crystal structure observed for the phosphate binding loop region (228–244), and the full-length *Cl*CRY4 with CTT (498–525) has not been crystalized. However, HDX-MS analysis effectively captures these structurally unresolved regions, revealing their dynamic and solvent-exposed nature.

**Figure 3.**
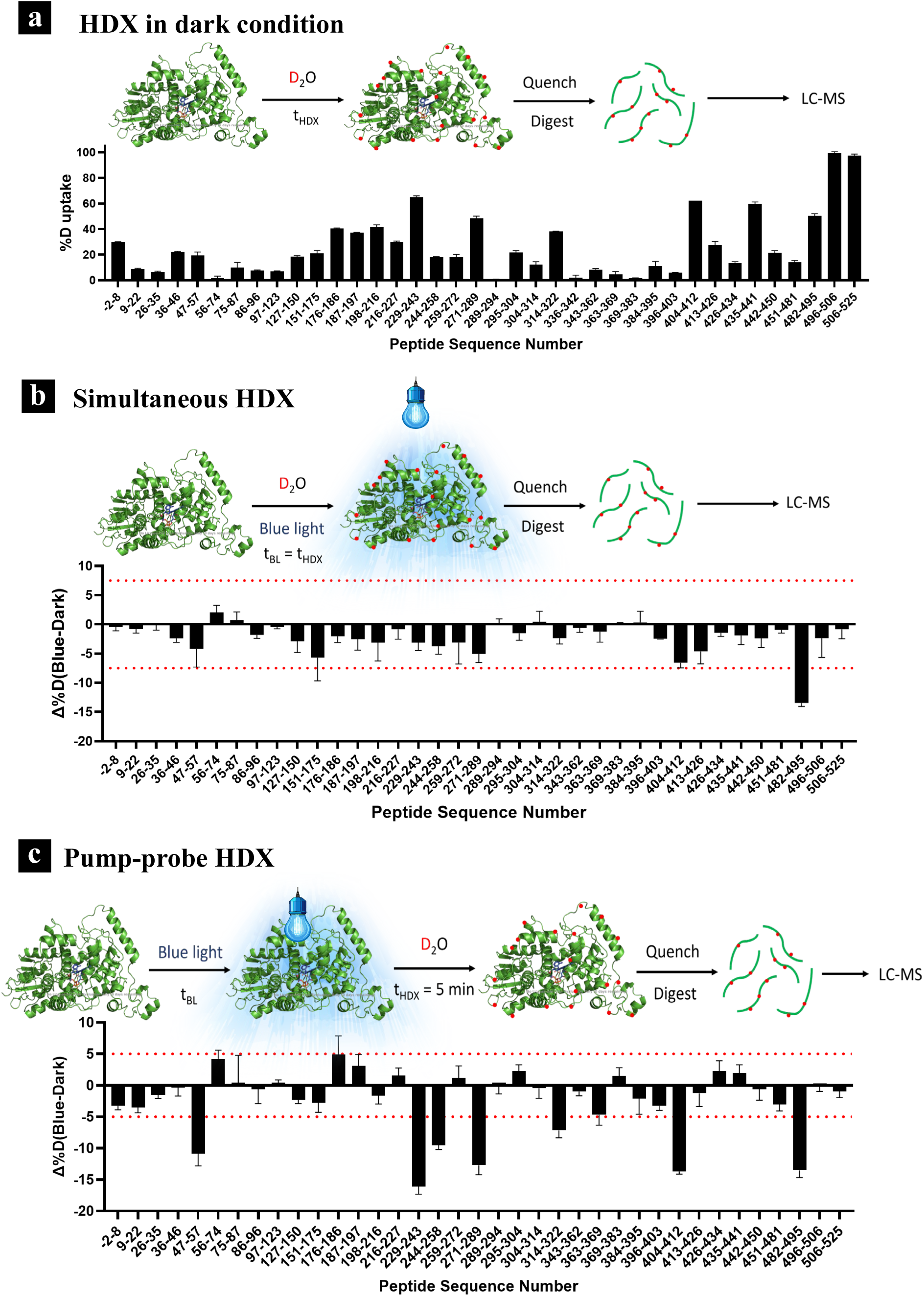
Hydrogen–deuterium exchange mass spectrometry (HDX-MS) of *Cl*CRY4 at 10 °C. **(a)** Per-peptide deuterium uptake (%D) after 5 min of exchange under dark conditions; the schematic above illustrates the dark HDX workflow (D2O labeling for tHDX, quench, pepsin digestion, LC-MS). **(b, c)** Light-induced change in deuterium uptake, Δ%D = %Dblue light − %Ddark, for each peptide measured by **(b)** the simultaneous HDX method (blue light and D2O labeling applied together; tBL = tHDX = 5 min) and **(c)** the pump–probe HDX method (5 min blue-light pump followed by a 5 min D2O probe in the dark). Negative Δ%D indicates light-induced protection. Dotted lines denote the ±2σ confidence interval; peptides are ordered by sequence position along the x-axis. Data are mean ± propagated error from 2 biological replicates.

#### Simultaneous HDX-MS Under Blue-Light Exposure

Initially, we investigated structural changes in *Cl*CRY4 induced by photoexcitation, under conditions where HDX-MS experiments were conducted under continuous blue-light exposure (450 nm, 250 lux), i.e., the time of deuterium exchange initiation matched the duration of light exposure. The experimental conditions were otherwise identical to those used in the dark-state measurements. Deuterium uptake profiles after blue-light exposure (blue traces) were compared to those under dark conditions (red traces) for all peptides (**Figures S5** for 25 °C and **S6** for 10 °C). The difference in deuterium uptake upon blue-light exposure (Δ(%D)) was calculated using the following equations:

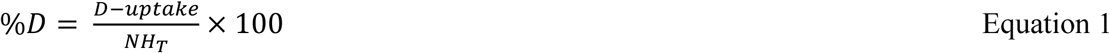

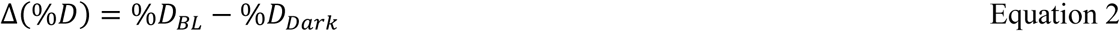

Where *NH_T_* are total exchangeable amides, %*D_BL_* and %*D_Dark_* are the percentage of D-uptake after light-activation and in the dark, respectively.

Peptides exhibiting significant changes in deuterium uptake (Δ(%*D*) > 1σ) over the 20 min of light-activation are mapped onto the protein structure in **Figure S7** for both temperatures. At 25 °C, a mixture of all three FAD redox states was present throughout the experiment, making it difficult to attribute structural changes to a single redox state, especially the signaling state FADH^•^. However, overall increased protection upon light activation was observed. Six peptides exhibited light induced changes close to or above the 1σ threshold. Among these, four peptides are associated with the FAD-binding region – 229-243 (part of the phosphate binding loop), 271-289 (protrusion motif), 384-395 (contains W395 or Trp1) and 396-403 (continuation of the W395-containing helix). The other two peptides constitute the C-terminal region just before the CTT - 464-481 and 481-495 (α-helix 22). At 10 °C, five peptides exhibited increased protection upon light activation. Four of these peptides (229-243, 271-289, 396-403, and 482-495) also showed protected at 25 °C. However, two peptides observed at 25 °C (384-395 and 464-481, both α-helices), did not show uptake in the dark at 10 °C and thus no detectable change upon light activation. Additionally, the peptide 404-412 (C-terminal lid, a loop structure) exhibited increased protection at 10 °C, though it was not observed to be significantly different at 25 °C. At higher temperature, this structurally flexible loop saturates in D-uptake, whereas at lower temperature its exchange remains in the growing range. Across both temperatures, structural changes were localized near the FAD-binding region and the C-terminal region that interacts with the Trp/Tyr end of the electron transfer chain.

At 10 °C, FADH^−^ formation was negligible before ∼8 minutes, with only small amounts accumulating between 10–20 minutes (**Figure 2c**). Therefore, structural changes observed between 10 seconds and ∼7 minutes can be predominantly attributed to FADH^•^, the putative signaling state. The Δ(%*D*) data for all peptides at 5 minutes of HDX at 10 °C are shown in **Figure 3b**. However, the observed differences are close to the experimental error threshold, with four peptides exhibiting changes close to 1σ and only one peptide (482–495) surpassing 2σ (dotted line). Given these limitations with the simultaneous HDX method, a modified experimental approach - **Pump-Probe HDX-MS -** was developed to better resolve light-induced structural dynamics in *Cl*CRY4.

#### Pump-Probe HDX-MS for Improved Detection of Light-Induced Structural Changes

To enhance the detection of light-induced structural changes in the putative signaling state of *Cl*CRY4, **Pump-Probe HDX-MS** approach was developed and performed at 10 °C. Based on the time-dependent deuterium uptake traces (**Figure S6**) and the slow reoxidation kinetics following blue-light exposure (**Figure S8**), a fixed HDX duration of 5 minutes was chosen as the “probe” phase. In this approach, *Cl*CRY4 is first “pumped” with blue light (450 nm, 250 lux) for defined durations (10 s, 20 s, 1.5 min, 3 min, 5 min, and 20 min), followed by 5 min of HDX probing conducted in the dark. These pump time points were chosen to allow direct comparison between HDX-MS and UV/Vis experimental protocols. The selected pump times account for: (i) initial short exposure (10 s and 20 s), (ii) half-life of FADH^•^ formation (1.5 min), (iii) peak FADH^•^ accumulation (3 min and 5 min), and (iv) an extended exposure (20 min) to access further structural changes from minor FADH^−^ formation. The probe time was chosen such that optimum deuterium uptake is achieved while minimizing reoxidation. All other experimental conditions remained consistent with the simultaneous HDX-MS protocol. By allowing light-induced structural rearrangements to occur before deuterium exchange, this approach improved the precision of conformational change detection at a given temperature.

**Figure S9** presents deuterium uptake traces for peptide 229–243 (phosphate binding loop), comparing simultaneous HDX-MS and pump-probe HDX-MS methods. Pump-probe HDX exhibited reduced experimental errors, leading to lower propagated errors in Δ(%*D*) calculations. More importantly, since deuterium exchange occurred after photoactivation, ground-state structural dynamics contributed less interference, enhancing precision in probing excited-state dynamics. As a result, pump-probe HDX is more sensitive to the detection of light-induced conformational changes compared to the simultaneous HDX.

The Δ(%*D*) values for all peptides across pump times (with a 5-minute probe phase) are summarized in **Figure 3c**, alongside comparisons with dark-state (%*D*) uptake and simultaneous HDX-MS results **(Figure 3a and b**). Most peptides exhibited reduced deuterium uptake upon blue-light exposure, with seven peptides exceeding the 2σ threshold (dotted lines), indicating light-induced structural protection in specific regions. With simultaneous HDX approach, four of these peptides (229-243, 271-289, 404-412, and 482-495) showed increased protection with lower confidence limits. Three additional peptides (47-57, 244-258, and 314-322) showed increased protection only in pump-probe HDX. Peptides 47-57 and 244-258 contain loops and small helices, with residues in the 244-258 (part of phosphate binding loop) interacting with FAD isoalloxazine ring and phosphate groups. Peptide 314-322 is a loop region containing key residues W318 and Y319, involved in the electron transfer chain.

A full comparison of all peptides showing D-uptake changes across methods (pump-probe HDX at 10 °C, simultaneous HDX at 10 °C, and simultaneous HDX at 25 °C) is provided in **Table S2**. Notably, peptide 396-403, which exhibited increased protection in simultaneous HDX at 10 °C, was not observed in pump-probe HDX. This is likely due to the 5-minute probe phase not capturing sufficient D-uptake, as the structural change in 396-403 was only detectable after 15 minutes in simultaneous HDX, where it had higher D-uptake in the dark-state.

#### Targeting C-terminal tail Peptides with Fast Probing

Because the C-terminal tail is the element most directly tied to cryptochrome function and signaling, we first asked whether the *Cl*CRY4 CTT participates in, and responds to, the light-driven photocycle, and on what timescale relative to accumulation of the FADH^•^ signaling state. The C-terminal tail (CTT) of cryptochromes and photolyases plays a crucial role in their function (32–34). Notably, deletion of the CTT in Drosophila CRY (*Dm*CRY) and in avian CRY4 abolishes magnetic sensing, whereas the blue-light photochemical response persists, possibly with minor alteration. HDX data at 10 °C revealed that the two peptides corresponding to the *Cl*CRY4 CTT (496-506 and 506-525) exhibited ∼80% deuterium uptake at the first time point (10 s) under dark conditions, reaching full saturation within 1 minute (**Figure 3a** and **S6**). Consequently, with the 5-minute probe time in pump-probe HDX, detection of blue-light induced changes in these rapidly exchanging peptides was challenging.

To specifically target these fast-exchanging CTT peptides, a 40-second deuterium exchange probing time was chosen, capturing D-uptake during the growing phase rather than at saturation. Under these conditions, both peptides exhibited a significant increase in structural protection upon blue-light exposure, starting from 10 seconds of illumination and persisting through 20 minutes of exposure (**Figure 4**). These results demonstrate that fast probing effectively captures light-induced conformational protection in the *Cl*CRY4 CTT region, highlighting its functional importance in photoactivation and signaling. Although this protection begins before substantial FADH^•^ accumulation, it persists throughout the period over which FADH^•^ builds up, and is therefore relevant to the signaling state. Having established that the CTT responds rapidly before the signaling state has fully formed, we next mapped the light-induced conformational changes across the remainder of the protein to define the full structural response and its relationship to FADH^•^ accumulation.

**Figure 4.**
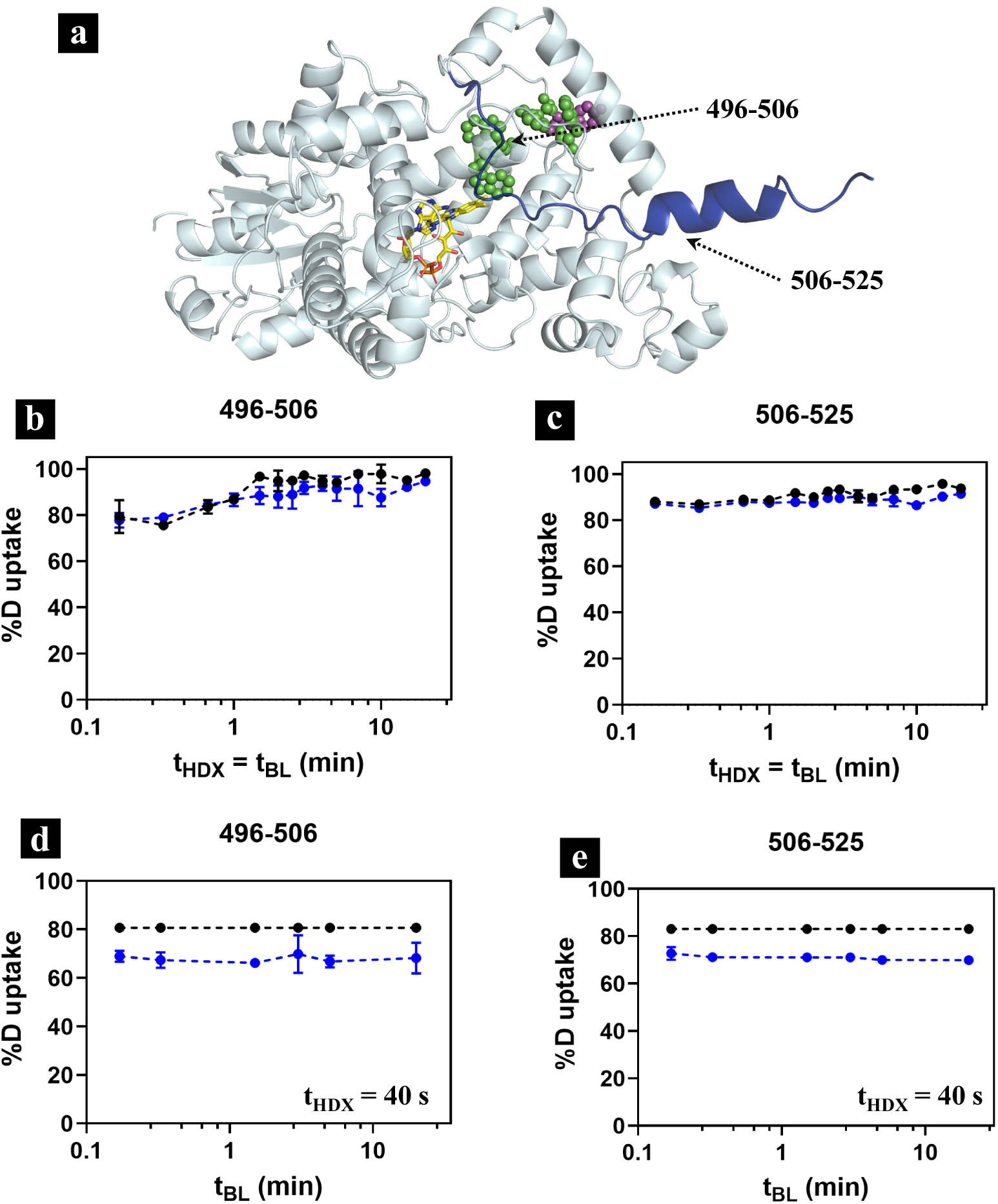
Light-induced structural protection in the C-terminal tail (CTT) of *Cl*CRY4 detected by pump-probe HDX-MS with fast probing at 10 °C. (a) AlphaFold model of *Cl*CRY4 with the CTT (residues 496–525) highlighted in blue; the FAD cofactor (yellow sticks), tryptophan tetrad (green spheres), and Y319 (magenta spheres) are shown for reference. **(b, c)** Simultaneous HDX-MS (tHDX = tBL) of CTT peptides 496–506 (b) and 506–525 (c) under dark (black) and blue-light (blue) conditions. Both peptides reach ∼80–90% deuterium uptake within the first time point (10 s), reflecting high solvent accessibility and conformational flexibility in the CTT. At these near-saturation exchange levels, light-induced protection is difficult to resolve. **(d, e)** Pump-probe HDX-MS with a fixed 40-s deuterium exchange probe of peptides 496–506 (d) and 506–525 (e) under dark (black) and blue-light (blue) conditions as a function of blue-light exposure time (tBL). Two changes enable detection of light-induced protection: first, the pump-probe design decouples photoactivation from deuterium exchange, allowing conformational effects to be sampled independently of illumination time; second, the short 40-s probe captures uptake during the growing (sub-saturation) phase of exchange, where the CTT retains sensitivity to changes in protection. Both peptides show significant light-induced protection from 10 s of illumination through 20 min.

### Signaling State Specific Structural Changes

The Δ(%D) plots and spatial mapping of seven peptides exhibiting significant changes upon blue-light excitation using the pump-probe HDX approach are shown in **Figure 5** (individual peptides in **Figure 5b–h**; mapped onto the *Cl*CRY4 structure in two orientations in **Figure 5a**, with FAD (yellow), the Trp tetrad (green and blue where light-activated change observed), and Y319 (blue) shown as sticks. The time intervals can be characterized based on FADH^•^ kinetics into three distinct zones: zone 1 (initial excitation - 10s and 20 s), which is the early response phase when FADH^•^ is ∼10% of the total FAD steady state concentration; zone 2 (half-life to complete formation – 90 s to 300 s), corresponding to the growth phase where FADH^•^ accumulation progresses; and zone 3 (extended exposure – 1200 s), a later-stage response that likely represents minor FADH-formation.

**Figure 5.**
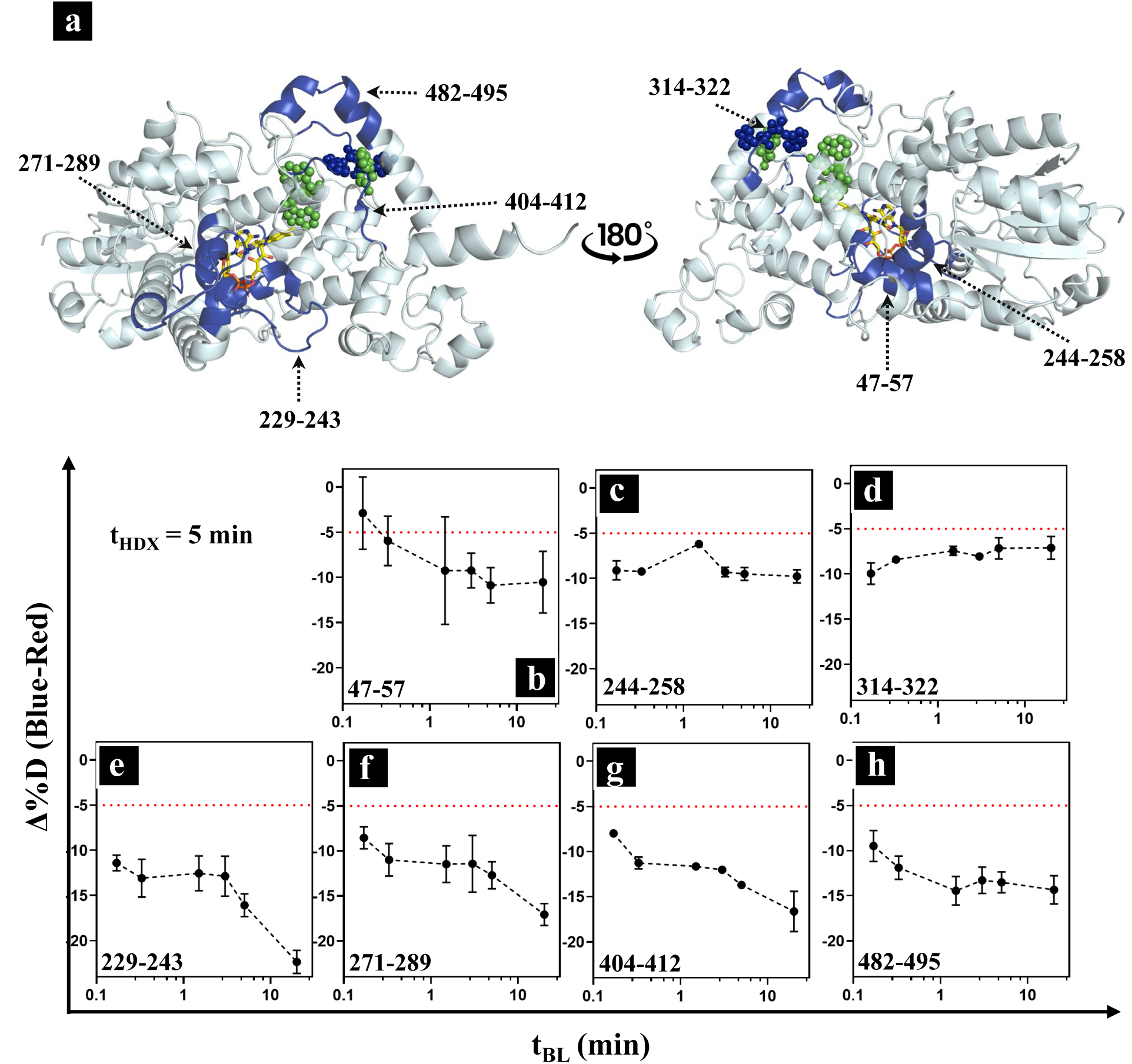
Seven peptides exhibit light-induced protection in *Cl*CRY4, resolved by pump-probe HDX-MS (10 °C, 5-min probe), revealing three kinetic classes of conformational response. (a) The seven significantly protected peptides (Δ%D beyond 2σ) mapped onto the *Cl*CRY4 structure in two orientations related by a 180° rotation (FAD, yellow; Trp tetrad, green; Y319, magenta). **(b–h)** Δ%D (blue-light − dark) versus blue-light pump time (tBL) for peptides (b) 47-57, (c) 244-258, (d) 314-322, (e) 229-243, (f) 271-289, (g) 404-412, and (h) 482-495. The dotted red line marks the −2σ significance threshold (≈ −5% Δ%D). The seven peptides segregate into three kinetic classes based on how their protection evolves relative to FADH• accumulation (Figure 2). **Class I** peptides (c: 244–258 and d: 314– 322) — both located at the electron-transfer chain — reach near-maximal protection by the earliest pump time (10 s), before substantial FADH• has accumulated, indicating an immediate structural response to radical-pair formation. **Class II** peptides (e: 229–243, f: 271–289, g: 404–412, h: 482–495) show early-onset protection that progressively deepens over the full 20 minutes, tracking the buildup of the FADH• signaling state. **Class III** (b: 47–57), the most distal peptide from the FAD pocket in the N-terminal αβ-domain, displays an onset and gradual deepening, consistent with a conformational change that propagates outward from the FAD-proximal core. Data are mean ± propagated error from 2 biological replicates.

These pump–probe time courses resolve the seven peptides into three kinetic classes defined by how their protection evolves relative to FADH^•^ accumulation. *Class I* comprises the two loops flanking the electron-transfer chain, peptide 244–258 (**Figure 5c**), which contacts the FAD isoalloxazine ring and phosphate groups, and peptide 314–322 (**Figure 5d**), which contains the chain residue W318 and the candidate fifth redox-active Y319. Both reach near-maximal protection at the earliest pump time (10 s, zone 1), when FADH^•^ is only ∼10% of the steady-state FAD pool and peptide 314–322 partially relaxes at longer exposures, indicating an immediate and partly transient response that tracks radical-pair formation rather than FADH^•^ build-up. *Class II* comprises peptide 229–243 (phosphate-binding loop; **Figure 5e**), peptide 271–289 (protrusion motif; **Figure 5f**), peptide 404–412 (C-terminal lid; **Figure 5g**), and peptide 482–495 (α22 helix; **Figure 5h**). These regions show significant protection by 10 s that deepens progressively across zones 1–3 (10 s to 20 min), paralleling the rise and accumulation of the FADH^•^ signaling state. *Class III* consists of peptide 47–57 (Figure 5b), the most distal region from the FAD pocket in the N-terminal αβ-domain and the only peptide lacking significant protection at 10 s; its protection develops gradually over the following minutes, consistent with a secondary conformational change that propagates outward from the FAD-proximal core. Together, the three classes describe a spatiotemporal hierarchy of light-induced protection that begins at the electron-transfer chain, expands through the FAD-proximal shell as the signaling state accumulates, and finally reaches the protein periphery.

These observations highlight that photoinduced structural rearrangements occur before full FADH^•^ accumulation and that certain regions undergo dynamic conformational shifts depending on the FAD oxidation kinetics.

Taken together, the spatial distribution of light-induced protection reveals a coherent structural picture: the two ends of the electron transfer chain tighten immediately upon photoexcitation while their surrounding shells progressively consolidate as the FADH• signaling state accumulates. As shown by fast probing in **Figure 4d** and **e**, the C-terminal tail also becomes rapidly protected, but to a small extent, indicative of a localized ordering rather than a global stiffening. We note that the mid-CTT structure of *Cl*CRY4 contains a conserved motif, Met504–Glu505–Met506, that can be modeled using alpha fold as undergoing close interactions with specific side chains within the body of *Cl*Cry4 that undergo protection in light; these branch out in opposite directions toward the FAD binding pocket and the terminal Trp (**Figure 9c**). In particular, the intramolecular contacts involving Glu505 place it at the hub of the observed light induced changes and can be tested below through the introduction of a conservative point mutation that perturbs the geometry of the side chain while preserving its charge (see below).

### Bimodal D-uptake Behavior Observed in the Phosphate Binding Loop Region

Under the HDX-MS conditions in this study, bimodal deuterium uptake behavior was observed in only one region of *Cl*CRY4 protein covered by peptide 229–243 and corresponding to the first half portion of the phosphate-binding loop (PBL – 234-251). This loop is unresolved in the available crystal structure (PDB 6PU0); however, the AlphaFold model of *Cl*CRY4 (AF-A0A386QUR4-F1-model_v4) places it in proximity to one of the phosphate groups of the FAD cofactor (**Figure S10**), suggesting a potential interaction in the modeled conformation. Closer analysis revealed that a truncated version of this peptide (residues 232–243) yielded more robust bimodal fits (*See SI* for detailed analysis and fit results). This refined peptide includes K234, positioned to form a direct interaction with the phosphate group in the AlphaFold-modeled conformation (Figure S10), as well as two conserved prolines critical to the loop structure. Consequently, bimodal analysis was conducted using peptide 232–243.

The double-binomial fitting of the isotopic envelopes was performed using HX-Express v3 (35), and details for each replicate are provided in the *Supporting Information*. **Figure 6** presents the result from two biological replicates, examining the evolution of bimodal behavior under two conditions: (1) the ground state (dark condition) as a function of deuterium exposure time, and (2) the photoexcited state (blue-light condition) from the pump-probe approach, where deuterium exchange time was fixed at 5 minutes and the time of blue-light exposure was varied.

**Figure 6.**
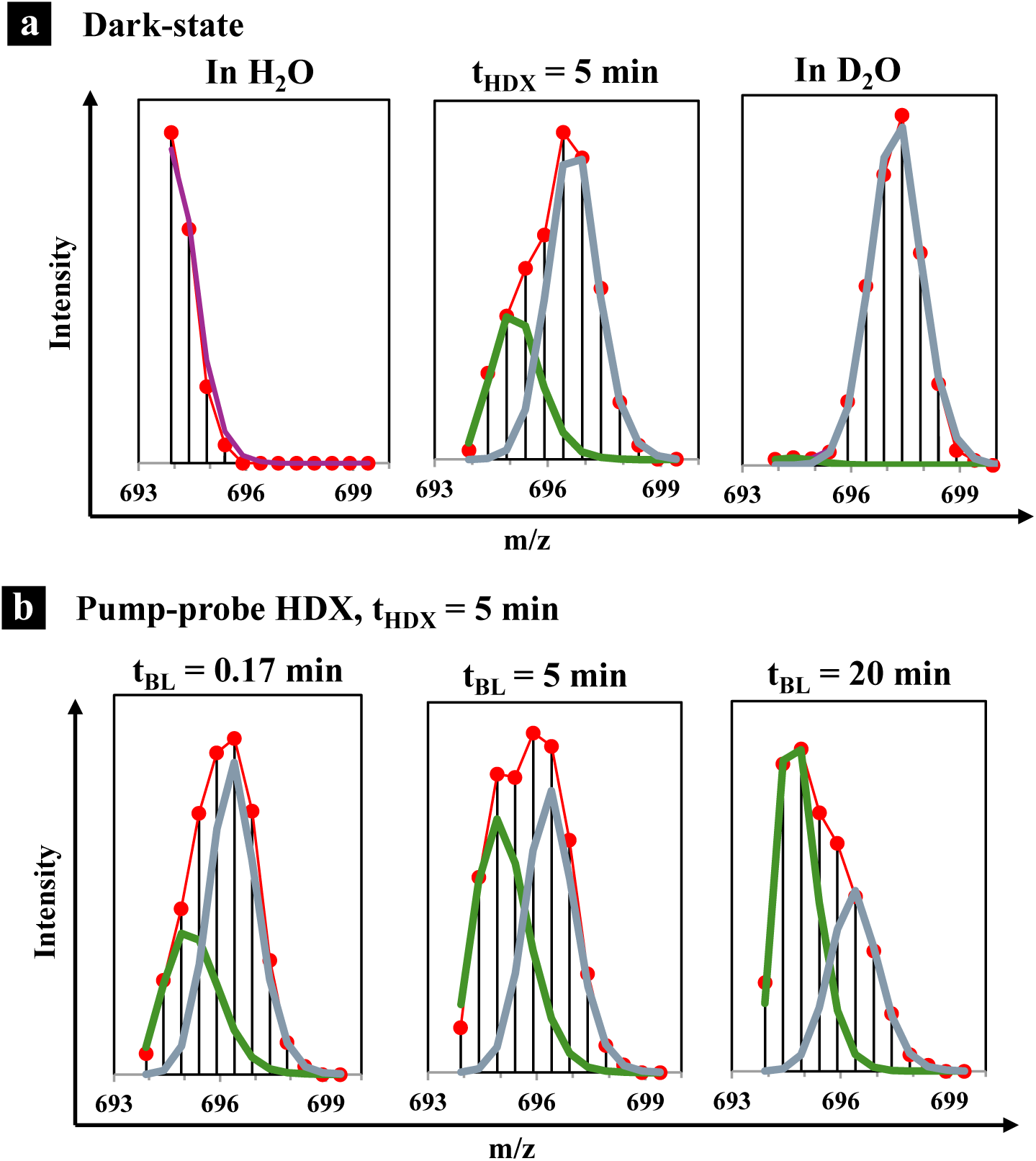
Bimodal isotopic envelopes of the phosphate-binding-loop peptide 232–243 resolved by double-binomial fitting. Mass spectra (red dots) of the +2 charge state of peptide 232–243 were fit using HX-Express v3 to resolve two coexisting conformational populations: a more protected conformer (lower m/z, green) and a more solvent-exposed conformer (higher m/z, grey). The composite fit is shown in red. **(a) Dark-state (FADox)** reference spectra at three exchange conditions: undeuterated control in H2O (left), 5 min of deuterium exchange at 10 °C (center), and a fully deuterated control obtained by overnight incubation in 100% D2O at 40 °C to quantify back-exchange during the work up (right). At tHDX = 5 min, both populations are clearly resolved, with gradual increases in deuterium incorporation relative to the undeuterated reference. **(b) Pump-probe HDX-MS** spectra at a fixed probe time of tHDX = 5 min, recorded after 0.17, 5, and 20 min of prior blue-light exposure (tBL) at 10 °C. The relative intensity progressively shifts toward the lower-m/z (more protected) population with increasing blue-light exposure, indicating a light-induced conformational redistribution that favors the more protected state. This behavior is consistent with EX2 exchange kinetics and implicates a photoinduced shift in the equilibrium position of the phosphate-binding loop, possibly via a proline *cis*–*trans* isomerization within two conserved prolyl bonds.

In each case, the centroid of the bimodal peaks is represented in the deuterium uptake of the two populations, while the relative intensities indicate their abundance. In the dark conditions (**Figure 6a**), corresponding to fully oxidized FAD, both populations exhibited a gradual increase in deuterium uptake with increasing exchange time. The relative population ratio remained stable during the first 2–3 minutes but began to shift toward the higher m/z population by 20 minutes, suggesting a relatively slow conformational transition. Note that the bimodal fits show a higher standard error as the two populations have significant overlap.

The observed bimodal exchange pattern is consistent with EX2 kinetics rather than the alternative EX1 mechanism. In classical EX1 kinetics, proteins undergo rapid, correlated unfolding events that result in highly deuterated subpopulations that grow over time. In contrast, the bimodal behavior observed for peptide 232-243 shows both populations gradually increasing D-uptake as a function of exchange time (**Figure S12**), characteristic of EX2 exchange where individual amides exchange independently according to local structural environment. This pattern implicates conformational heterogeneity within the phosphate-binding loop, where the peptide samples two distinct structural states with different degrees of solvent accessibility and hydrogen bonding patterns. The persistence of both populations throughout the time course, combined with their parallel increases in deuterium incorporation, strongly supports the interpretation that this region exists in a dynamic equilibrium between two conformationally distinct substates rather than undergoing global cooperative unfolding.

During the light-induced changes through pump-probe experiments (**Figure 6b**), in contrast to the evolution of bimodal behavior in the dark, the deuterium uptake of both populations remained largely unchanged over varying durations of blue-light exposure. However, the relative intensity of the lower m/z population increased progressively with longer blue-light exposure times, indicating a light-induced conformational shift that favors the population that exhibits more protection.

### Testing the Role of E505 in Light Induced Signaling

The patterns of HDX protection within *Cl*CRY4 upon blue light illumination, together with the identification of a conserved motif Met504–Glu505–Met506 midway along the CTT, leads to a working molecular model where Glu505 acts as a “clamp” in creating the light induced conformational changes essential for signaling. As shown in **Figure 9c**, the carboxylate of E505 can be modeled proximal to residues lining the FAD-proximal cleft and the protrusion/phosphate-binding regions (e.g., F404, H406, R409, P235, Q287, V282) within regions shown to undergo increased HDX protection in the presence of light. We therefore generated the conservative point mutant E505D, that preserves negative charge but shortens the side chain by one methylene unit, thereby capable of perturbing the local geometry of contacts while leaving the overall fold and the FAD-binding architecture intact. Comparison of E505D with wild-type (*Cl*CRY4 (WT) was carried out across three readout protocols: steady-state photochemistry, dark-state HDX, and pump–probe HDX.

The photochemistry of E505D was markedly altered (**Figure S17a**). At 10 °C, photoreduction of FAD_ox_ was slowed and remained incomplete: rather than the near-complete conversion seen for WT, FAD_ox_ in E505D never fully depleted and retained roughly 40% of its initial population over the 20 min illumination. Correspondingly, accumulation of the FADH^•^ signaling state plateaued at only ∼50–60%, compared with the substantially higher levels reached by WT before the slower decay associated with FADH^−^ formation. In the dark, moreover, reoxidation of E505D was faster than that of WT measured under the same conditions (10 °C), with FAD_ox_ recovering more rapidly to its ground-state level. Together, these observations indicate that the E505D substitution both impedes forward progression into the signaling state and destabilizes the photoreduced state once formed, consistent with a loss of the stabilizing contacts that E505 normally provides.

Dark-state HDX-MS also reveals that the E505D ground state is intrinsically more solvent-exposed than that of WT (**Figure S17b**). Across the nine peptides that report on the light-responsive regions, three show slightly elevated dark deuterium uptake in the mutant relative to WT: the phosphate-binding loop (229–243), the protrusion motif (271–289), and the C-terminal lid (404–412). The CTT peptides (496–506 and 506–525) were already near-saturated in both proteins and thus uninformative in the dark, while the remaining peptides were largely unchanged. The increased baseline exchange in 229–243, 271–289, and 404–412 indicates that disrupting the E505 modestly loosens the regions of the FAD-proximal shell that are seen in WT to undergo significant protection upon illumination (**Figure 5**).

The decisive test comes from pump–probe HDX-MS, which quantifies the change in protection between the dark ground state and the blue-light-activated state for each variant. Because the comparison is made internally—each protein’s light state referenced to its own dark state as Δ%D (blue−dark)—it is a robust indicator of the different absolute redox compositions of the two proteins. In WT, all seven peptides show significant light-induced protection organized into the three kinetic classes described above (**Figure 5**). In E505D, this response is largely abolished (**Figure 7b–h**): the Class I electron-transfer-chain loops (244–258, 314–322) and the Class III distal peptide (47–57) no longer show the WT protection, and the protrusion and C-terminal-lid peptides (271–289, 404–412) likewise lose their light-induced protection. Only the α22-helix peptide 482–495 retains a WT-like protected response (**Figure 7h**), and the phosphate-binding loop 229–243 recovers only a partial, delayed protection at the longest blue-light exposures (**Figure 7e**). The selective survival of the 482–495 response is informative: it serves as an internal control showing that the mutation does not globally eliminate the protein’s capacity for light-induced structural change, but instead specifically disrupts the clamp-coupled regions, underscoring that the lost responses reflect a targeted breakdown of the clamp network rather than a non-specific destabilization.

**Figure 7.**
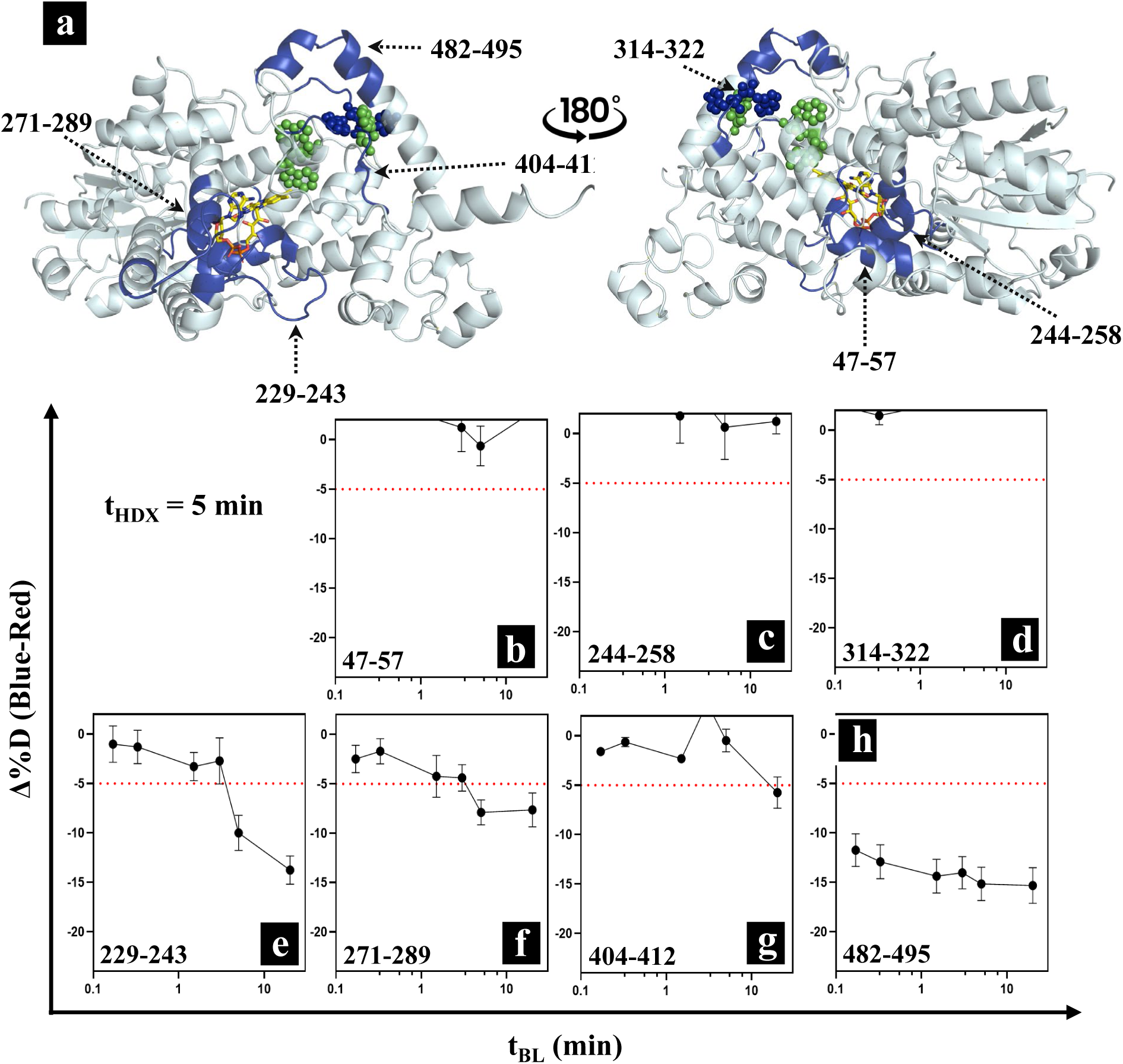
Pump–probe HDX-MS of the E505D variant compared with wild-type *Cl*CRY4. (a) The seven light-responsive peptides mapped onto the *Cl*CRY4 structure in two orientations (rotated 180°); the regions are identical to those in Figure 5a (FAD, yellow; phosphate-binding loop and adjacent elements, blue; space-filling Trp tetrad, green). (b–h) Change in deuterium uptake upon blue-light activation, Δ%D (blue−dark), versus blue-light pump time (tBL) at a fixed 5-min probe, 10 °C, for peptides 47–57 (b), 244–258 (c), 314–322 (d), 229–243 (e), 271–289 (f), 404–412 (g), and 482–495 (h). The red dotted line marks the −2σ significance threshold (≈ −5% Δ%D). In wild type (Figure 5) all seven peptides show significant light-induced protection; in E505D this response is largely abolished except for peptide 482–495 (h), which retains a wild-type-like response, with only partial, delayed protection recovered in 229– 243 (e) and possibly 271-289 (f).

Interpreting these differences requires accounting for the fact that E505D accumulates FADH^•^ signaling state but at a ca. 50-60% level of the WT. In principle, a reduced Δ%D in the mutant could therefore arise either because less signaling state is present or because the per-molecule conformational response is genuinely disrupted. Two observations argue that the effect is a real disruption of the clamp rather than a simple consequence of reduced signaling-state population. First, the dark ground state of E505D is already looser in the FAD-proximal shell (229–243, 271–289, 404–412), indicating that the E505 contacts shape the resting conformation independently of how much signaling state forms. Second, light-induced HDX protection is lost even in regions where signaling state does accumulate, and occurs in a spatially selective pattern (sparing 482– 495) rather than uniformly attenuated, which is the signature expected of a broken local contact network rather than a uniform dilution of the response. We therefore interpret the E505D data as evidence that E505 is functionally required for the light-induced clamp, while noting explicitly that the lower signaling-state yield of the mutant precludes a strictly quantitative, magnitude-for-magnitude comparison of Δ%D between the two proteins.

## Discussion

### Temperature-Dependent Photochemistry and Mechanistic Insights

Steady-state UV-vis absorption spectroscopy (Figure 2) demonstrates that *Cl*CRY4 photochemistry is strongly temperature-dependent. Lower temperatures slow the reduction of FAD_ox_ and the formation and decay of FADH^•^, with FADH^•^ persisting longer at 10 °C and 5 °C. These results are consistent with previous reports that temperature modulates cryptochrome photochemistry, possibly through effects on protonation equilibria or protein dynamics (19,36). However, the lack of a kinetic isotope effect in D_2_O (Figure S2) argues that proton transfer is not the rate-limiting step in FADH^•^ formation. Instead, the data support a model in which a conformational change or isomerization limits the accumulation of the signaling state. This contrasts with photolyases, where proton transfer from conserved residues is often rate-limiting and sensitive to solvent isotope substitution (37,38). In *Arabidopsis* CRY1, temperature also modulates the lifetime of the signaling state, but the effect is less pronounced than in *Cl*CRY4 (36). Conversely, *Drosophila* CRY has been reported to exhibit temperature-insensitive CTT release (39), suggesting that avian CRY4s may have evolved unique temperature-coupling mechanisms, potentially as an adaptation for environmental sensing during migration (19).

### Light-Induced Conformational Changes Revealed by HDX-MS

HDX-MS analysis reveals that blue-light exposure triggers localized structural protection in *Cl*CRY4, particularly in regions proximal to the FAD-binding site and the C-terminal region. The pump-probe HDX-MS approach (**Figure 3**), which decouples photoactivation from deuterium exchange, has led to improved precision and sensitivity (at a 2-σ level) for detecting these conformational changes. Seven peptides (**Figure 5**), including those in the phosphate binding loop, protrusion motif, C-terminal lid, and α22 helix, exhibit significant light-induced protection, indicating that these regions undergo conformational rearrangements upon photoexcitation. Notably, many of these changes occur rapidly, within 10 seconds of blue-light exposure, preceding the accumulation of the imputed signaling species FADH^•^. Specifically, two regions around the two sides of the electron transfer chain, one covering half of the phosphate binding loop (peptide 244-258) and another loop containing Tyr319 (peptide 314-322) show the highest change at the first experimental timepoint of 10 s, indicating structural changes that take place prior to the full accumulation of FADH^•^. Other regions exhibiting conformational changes show increasing protection over the 20 minutes of the experimental readout. This suggests that structural transitions are initiated early in the photochemical cycle, potentially priming the protein for downstream signaling. These results are consistent with recent studies on pigeon CRY4, where light-induced conformational changes were mapped to the phosphate binding loop and CTT regions using limited proteolysis and molecular dynamics (15,17,40). In *Drosophila* CRY, light-induced CTT release is well established (10,41), but the process is slower and dependent on subsequent phosphorylation events (39). Our observation that *Cl*CRY4’s CTT becomes rapidly protected upon light exposure suggests a direct and immediate conformational response, more akin to plant CRYs (42) than to insect CRYs.

The pump–probe HDX-MS data provide structural support for this interpretation: protection at the two electron-transfer-chain loops (peptides 244–258 and 314–322) is essentially complete within 10 s of illumination, well before substantial FADH^•^ accumulates; demonstrating that a structural rearrangement near the FAD pocket precedes the proton-uptake event inferred from steady-state kinetics. This is also true for FADH^•^ accumulation that appears identical in light and heavy water.

Viewed together, the pump–probe time courses (**Figure 5**) segregate the seven light-responsive peptides into three kinetic classes that map onto distinct stages of the photocycle: *Class I* (244– 258 and 314–322), located on either side of the electron-transfer chain, reaches near-maximal protection at the earliest pump time and tracks radical-pair formation rather than FADH^•^ accumulation. *Class II* (229–243, 271–289, 404–412, 482–495) shows early-onset protection that deepens progressively over 20 min, paralleling the rise of the FADH^•^ signaling-state population. *Class III* (47–57), the most distal peptide from the FAD pocket in the N-terminal αβ-domain, is the sole peptide without significant protection at 10 s and develops gradually, consistent with a secondary conformational change propagated outward from the FAD-proximal core. This spatiotemporal hierarchy, an instantaneous response at the electron transfer chain, followed by signaling-state-coupled tightening of the FAD-proximal shell, and delayed propagation to the protein periphery; directly links the photochemical kinetics of **Figure 2** to the structural dynamics and is reminiscent of the distal, millisecond conformational changes recently resolved by time-resolved crystallography in the animal-like cryptochrome *Cra*CRY (45). Crucially, this resolution into kinetic classes is enabled by the pump–probe HDX-MS strategy itself: by decoupling photoactivation from deuterium labeling, the method reports on the conformational state of the protein as a function of illumination time rather than averaging over the exchange window, allowing structural events that precede, accompany, and follow signaling-state accumulation to be separated. A comparable blue-light-induced, peptide-resolved conformational change in the C-terminal tail of *Drosophila* cryptochrome was recently characterized by HDX-MS (46), underscoring the general power of this approach for mapping cryptochrome signaling states; the present work extends it to a temperature-resolved, time-resolved pump–probe format that captures the *Cl*CRY4 activation pathway across its full kinetic range.

### Functional Role of the C-terminal Tail

The C-terminal tail (CTT) of *Cl*CRY4 displays rapid deuterium exchange and high solvent exposure in the dark, indicating a flexible and dynamic region (**Figure 3**). Using fast-probing HDX-MS (**Figure 4b**) has, however, made it possible to show that blue-light exposure induces protection in CTT. This observation is consistent with the observation that CTT truncation from the CRY4 of migratory birds abolishes magnetic sensing (19) and supports the hypothesis from limited proteolysis data that light will stabilize this region of protein (40). In *Drosophila* CRY, CTT release is required for interaction with the E3 ligase Jetlag and subsequent proteasomal degradation (10,48). Our findings of rapid CTT protection in *Cl*CRY4 indicate a local ordering of the tail, in opposition to the canonical response in *Drosophila* cryptochrome (*Dm*CRY) where photoactivation increases its solvent exposure to enable downstream interactions (10,41,46). That the *Cl*CRY4 CTT instead becomes more structured and protected upon illumination points to a distinct signaling logic, in which the activated tail becomes stabilized and oriented, rather than liberated, for productive partner engagement. This gain in protection is, moreover, detectable within the first 10 s of illumination, before the FADH^•^ signaling state has substantially accumulated, indicating that the CTT is not a passive downstream reporter but an early participant in the photocycle and that its ordering is among the first structural consequences of radical-pair formation rather than a late event contingent on full signaling-state population. Although the direction of the CTT response is thus opposite to that of *Dm*CRY, the finding that both tails respond rapidly and reversibly to blue light (recently resolved at peptide resolution for *Dm*CRY by HDX-MS (46)) supports the view that light-driven CTT mobilization, whether by release or by ordering, is a contributing early step in cryptochrome signal transduction.

### Bimodal Conformational Behavior in the Phosphate Binding Loop

One striking observation from this study is the bimodal distribution of deuterium uptake within the phosphate binding loop at peptide 229-243, observed to be unstructured in X-ray structures in the absence of blue light activation. The light induced impact on HDX-MS indicates a shift toward a more protected/folded conformer on the time frame of FADH^•^ production (**Figure 6**).

Notably, the phosphate binding loop contains Lys234, that is proposed to interact with the phosphate side chain of FAD within the folded conformer(49), as well as two proline residues, raising the possibility that *cis-trans* isomerization of these prolyl bonds underlies the observed bimodal HDX-MS behavior. Proline isomerization has been shown to introduce large protein structural changes that modulate protein function and signaling in numerous systems (e.g. 50,51). In folded proteins, the energetic barrier for such isomerization is high, making the process intrinsically slow (52), The uncatalyzed isomerization of proline typically occurs on the order of minutes at low temperatures such as 10 °C (42,50,53,54), a time frame closely matching the timescale of the bimodal HDX-MS changes observed in this study. We note that a coupling between light-induced structural changes and proline isomerization has been previously observed in other light-sensing proteins (43,44,52).

While the phenomenon of photoinduced conformational switching of a surface loop has not been reported in avian CRY4, a similar conformational heterogeneity was observed in the “serine loop” of mammalian CRY2, where loop dynamics modulate interaction with CLOCK: BMAL1 (55). In photolyases, the phosphate-binding loop is critical for FAD binding and repair activity but does not typically display such pronounced light-dependent switching (38).

### A Spatiotemporal Model Connects Quantum and Classical Processes in *Cl*CRY4 Magnetoreception

An integration of our HDX-MS findings and temperature dependent steady-state photochemical kinetics with literature findings (4,17,40,45) leads to a comprehensive model for *Cl*CRY4 activation that integrates quantum electron transfer processes with classical conformational dynamics (**Figure 8**). Upon blue-light excitation, FAD undergoes rapid photoreduction (picosecond timescale) to form the initial FAD^•-^ radical anion; this occurs concomitant with a sequential electron transfer cascade along the conserved tryptophan tetrad (W1→W2→W3→W4), possibly extending toward the terminal tyrosine residue (Y319). The formation of W4^•+^ occurs within nanoseconds, followed by rapid deprotonation (pKa ∼4.5-5.5) that may activate subsequent proton coupled electron transfer from Y319 to Trp^•^ on a sub-microsecond timescale.

**Figure 8.**
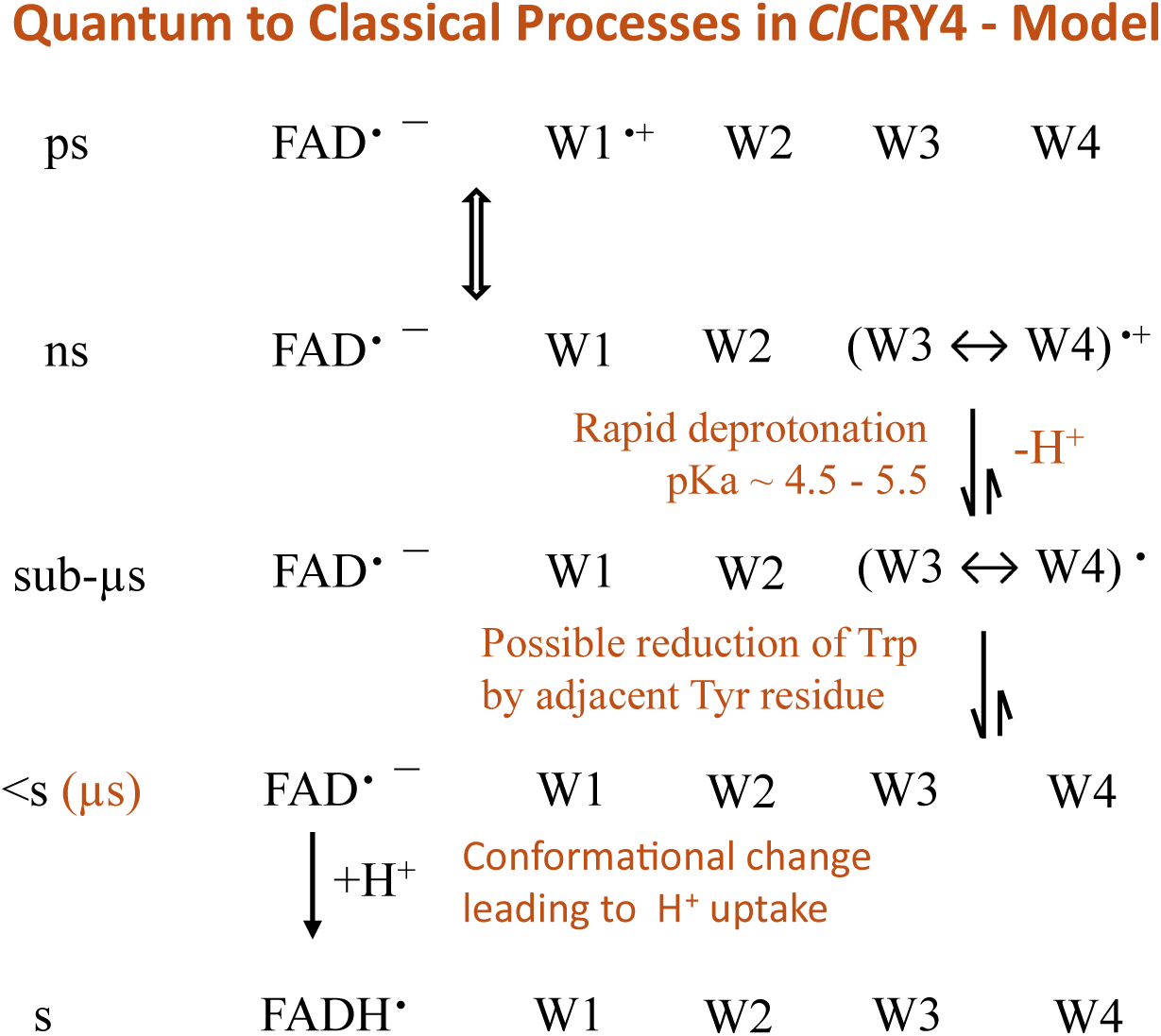
Proposed comprehensive model coupling quantum electron transfer to classical conformational change in *Cl*CRY4. Approximate timescales are shown on the left. Blue-light excitation generates FAD•– and a tryptophan cation radical (W1•+) within picoseconds; sequential electron transfer along the tetrad (W1→W2→W3→W4) produces the (W3 W4)•+ radical on the nanosecond timescale. Rapid deprotonation of the terminal tryptophan (pKa ≈ 4.5–5.5; −H+) and charge neutralization yield the neutral (W3 W4)• radical on the sub-microsecond timescale. Reaction of this state with an adjacent tyrosine may extend the free radical center further along the electron chain to produce a fully reduced Trp tetrad. A conformational change impacting the FAD pocket is coupled to proton uptake and formation of the neutral semiquinone FADH• (signaling state) on the second to minute timescale.

**Figure 9.**
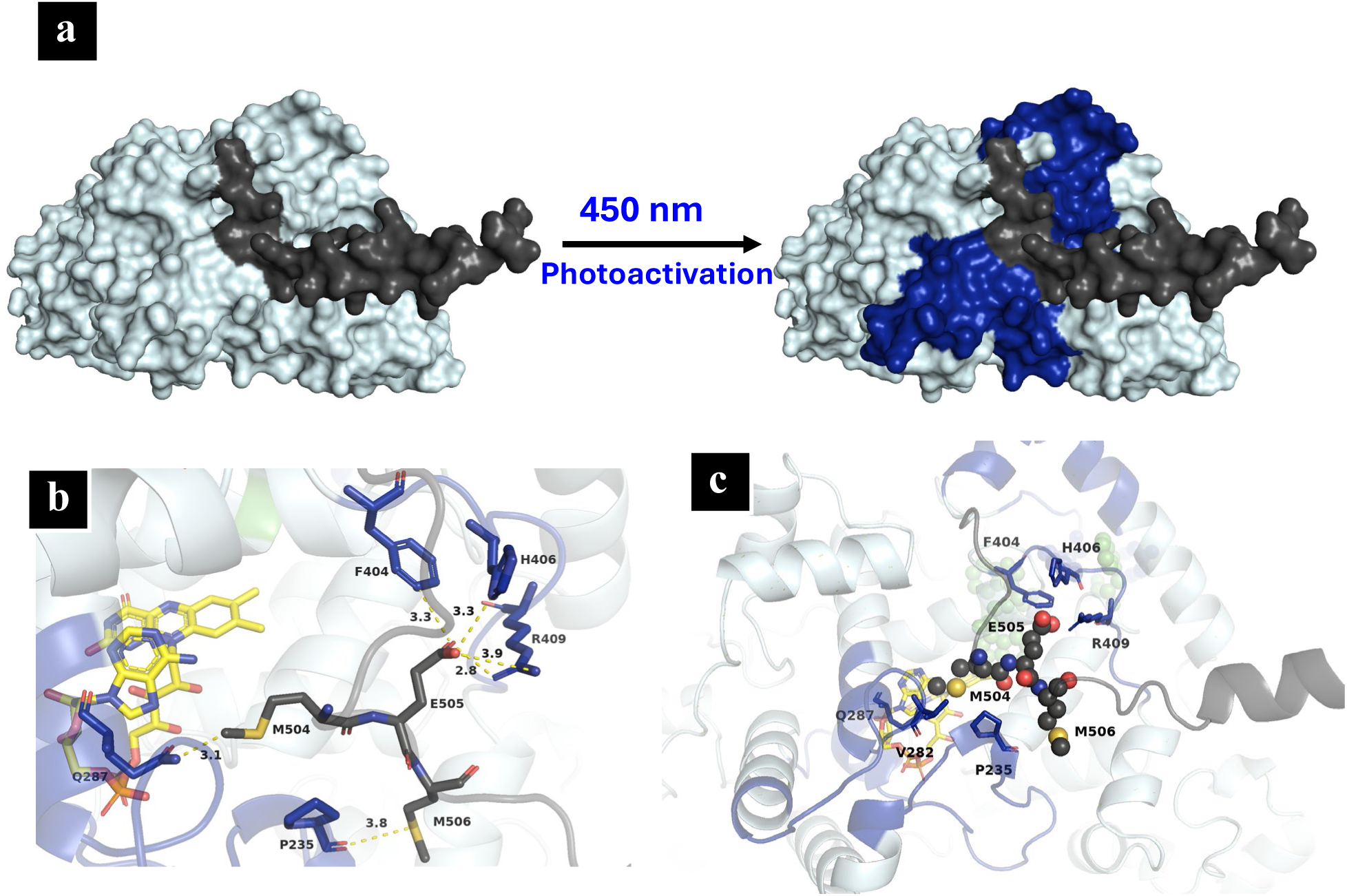
Photoactivation-induced conformational changes proposed to underlie downstream signaling in *Cl*CRY4. **(a)** Surface representations of *Cl*CRY4 in the dark (left) and after 450 nm photoactivation (right). Regions that gain protection upon illumination are blue; the CTT missing in the crystal structure is dark gray. **(b)** Close-up of the mid-CTT M504–E505–M506 motif and its hydrogen-bonding/contact network with protected core regions (e.g., F404, H406, R409, P235, Q287); selected distances (Å) are indicated. **(c)** Cartoon view of the same interaction network in the context of the full protein.

The generated radical sites are, thus, undergoing dynamical changes in photoinduced electron density/charge on a time scale faster than μs. From the pump-probe HDX-MS experiments that are collected on the time scale of seconds to minutes, these radical sites are seen to already be protected from HDX at the earliest time point (10 s) of photoactivation, a time where there is little accumulation of FADH^•^ (**Figure 5**). In the same time frame, some fraction of the C-terminal tail also becomes protected (**Figure 4**). The latter involves only a small component of the full length of the CTT, leading us to propose the motif Met504-Glu505-Met506 as the “glue” within the CTT that organizes the key internal regions of protein for subsequent steps (see below).

The generation of FADH^•^ from FAD^•-^ occurs with a half time of ca. 2-3 minutes (**Figure 2**) and is unperturbed by solvent D_2_O as noted above, implying a rate limiting conformation change that gates FADH^•^ formation on a time scale that is 10^6^-fold slower than the initial radical pair generation. The involvement of such a slow conformational change is notable in the context of the ability of *Cl*CRY4 to sustain reversible radical pair formation for a long enough time frame to establish and maintain magnetic sensitivity. As suggested by **Figure 8**, a sub-microsecond deprotonation of Trp (and possible involvement of an adjacent Tyr) will be the first step(s) ratcheting down the time scale of events, facilitated by the fact that both processes occur near a solvent interface with intrinsic reversibility based on known pKa values and redox potentials (56–58). Turning to the process of the conversion of FAD^•-^ to FADH^•^, this occurs within the buried core of the CRY4 structure and will be favored by an intrinsic pKa for unbound FADH^•^ of 8.3 to 8.5 (29,59) that may well be significantly elevated within the CRY4 active site. At this stage of FADH. formation, the door to reversible photoexcitation becomes effectively closed, creating the complete uncoupling of quantum behavior from classical protein restructuring and cell signaling.

### Unique Structural Features of the CTT in *Cl*CRY4

To visualize the spatial extent of these light-induced changes across the intact protein, **Figure 9a** maps the regions gaining HDX protection upon illumination (blue) onto the *Cl*CRY4 surface in the dark and light-activated states. Using AlphaFold to model the unstructured CTT of *Cl*CRY4, its mid-section, Met504-Glu505-Met506, 504–506, is found to reside in van der Waals distance to key side chains located within proximal regions of protein that undergo time dependent immobilization in blue light (**Figure 9b**). An emergent property from such a clamp-like configuration may be a positioning of the remaining length of the CTT for optimal interaction with downstream partnering and signaling proteins.

Support for the proposed model comes from the conservative E505D variant, in which the central mid-CTT residue is shortened by a single methylene while its charge is preserved. If E505 functions as a structural anchor that tethers the mid-CTT against the protein body, then perturbing its contacts should both loosen the resting state and impair the light-induced clamp - precisely what is observed. E505D begins from a somewhat less structured ground state that increases the dark-state deuterium uptake in the FAD-proximal shell (peptides 229–243, 271–289, and 404–412; **Figure S17**). Its photochemistry is correspondingly compromised, with slowed, incomplete photoreduction, a FADH^•^ signaling-state plateau of only ∼50–60%, and faster dark reoxidation, indicating that the mutation destabilizes the photoactivated state. Most tellingly, pump–probe HDX-MS shows that the light-induced protection that defines the WT response is largely abolished in E505D (**Figure 7**), with the notable exception of the α22-helix peptide 482–495, which retains a WT-like response and thereby demonstrates that the mutation disrupts the clamp network specifically rather than disabling light-induced structural change in general.

These results validate the central premise of the model: that the mid-CTT Met-Glu-Met motif, and E505 in particular, mediate the light-dependent conformational tightening that we propose couples photochemistry to downstream signaling. Such a clamp-like mechanism is distinct from the undocking or full release of CTT observed in other cryptochromes and may represent an evolutionary adaptation in birds to couple light-induced electron transfer with precise conformational control for environmental sensing. Future studies aimed at identifying CTT-interacting partners and characterizing these protein-protein interactions will be essential for elucidating the downstream signaling pathways that underlie magnetoreception in migratory birds.

### Broader Implications and Future Directions

In summary, this study establishes a framework for dissecting the molecular mechanism of cryptochrome activation, with implications for understanding its role in light sensing and signaling in biological systems. Collectively, the results indicate that *Cl*CRY4 integrates photolyase-like electron transfer with cryptochrome-specific conformational dynamics, supporting a hybrid mechanism for light sensing and signaling. The rapid, light-induced protection of the CTT and phosphate-binding loop, coupled with pronounced temperature sensitivity, distinguishes *Cl*CRY4 from both plant and insect CRYs, and may reflect evolutionary adaptation for magnetoreception in migratory birds. These findings provide a foundation for future studies to dissect the molecular determinants of CRY4 signaling, including mutational analysis of the CTT and loop regions, and investigation of protein-protein interactions with putative magnetoreceptor partners such as G protein coupled receptors (GPCRs), candidates for which have been identified in the avian retina (24), and MagR (25). Building directly on the model established here, several complementary experiments would extend and reinforce the conclusions. The proline-gating hypothesis for the phosphate-binding loop could be further substantiated by site-specific substitution of the two conserved prolines (P→A or P→Xaa), proposed to modulate the bimodal deuterium-uptake behavior of peptide 232–243. The CTT clamp could be examined in greater detail using a CTT-truncated construct, providing an independent structural correlate for the loss of magnetic sensing reported upon CTT deletion. Building on the clamp validation provided by E505D, further point substitutions within the Met-Glu-Met motif and its contact partners would map the determinants of the clamp at single-residue resolution and test whether signaling-state yield and conformational tightening can be tuned independently. Finally, extending the pump–probe HDX-MS framework developed in this work to measurements under an applied magnetic field, or in the presence of candidate signaling partners such as the cone-specific G protein and GPCRs noted above, represents a natural next step toward linking the conformational signaling pathway delineated here to field-dependent function *in vivo*. Limitations of this study include the temporal resolution restrictions (>10 s) of the current HDX-MS and the need for *in vivo* validation of the observed conformational changes. Time-resolved crystallography and single-molecule spectroscopy, as applied to Drosophila CRY, could further elucidate the structural transitions underlying *Cl*CRY4 activation (41). Ultimately, integrating these mechanistic insights with behavioral and physiological studies in birds will be essential to fully understand the role of CRY4 in magnetoreception and environmental adaptation.

## Acknowledgements

We are grateful to Susan Miller (UCSF) and Nigel G. J. Richards (Cardiff University) for their insightful input and critical discussion during group meetings. Kotchakorn T. Sriwong, Sabyasachi Sarkar, and Yaoyukun Jiang provided valuable feedback and thoughtful discussion of the HDX-MS data during lab meetings. We also acknowledge Paul M. Champion (Northeastern University), Adam R. Offenbacher (East Carolina University), and Michael C. Thompson (UC Merced) for stimulating discussions of this work during joint meetings. We thank the QB3/Chemistry Mass Spectrometry Facility at UC Berkeley for the use of mass spectrometry instrumentation. This work was supported by the National Science Foundation (grant no. 2231081, to J.P.K.) and the National Institute of General Medical Sciences (grant no. 1S10OD020062-01, to A.T.I.).

## Note added in deposition

While our manuscript was in the final stages of preparation, a related study reporting an HDX-MS investigation of cryptochrome 4 from the European robin (*Er*Cry4a) appeared (60). That work employs related but different methodologies and leads to a combination of similar and diverging implications from those presented here.

